# N-terminal ubiquitination of amyloidogenic proteins triggers removal of their oligomers by the proteasome holoenzyme

**DOI:** 10.1101/653287

**Authors:** Yu Ye, David Klenerman, Daniel Finley

## Abstract

Aggregation of amyloidogenic proteins is an abnormal biological process implicated in neurodegenerative disorders. While the aggregation process of amyloid-forming proteins has been studied extensively, the mechanism of aggregate removal is poorly understood. We recently demonstrated that proteasomes could fragment filamentous aggregates into smaller entities, restricting aggregate size[1]. Here, we show *in vitro* that UBE2W can modify the N-terminus of both αS and tau^K18^ with a single ubiquitin moiety. We demonstrate that an engineered N-terminal Ub modification changes the aggregation process of both proteins, resulting in the formation of structurally distinct aggregates. Single-molecule approaches further reveal that the proteasome can target soluble oligomers assembled from Ub-modified proteins independent of its peptidase activity, consistent with our recently reported fibril-fragmenting activity. Based on these results, we propose that proteasomes are able to target oligomers assembled from N-terminally ubiquitinated proteins. Our data suggest a possible disassembly mechanism by which N-terminal ubiquitination and the proteasome may together impede aggregate formation.

## Introduction

The 26S proteasome holoenzyme is responsible for selective protein degradation in eukaryotic cells[2]. Proteins selected for degradation are often covalently modified with ubiquitin (Ub) moieties, which are recognized by the proteasome[3]. The proteolytic activity required for degradation is provided by the 20S core particle (CP) of the holoenzyme, while the 19S regulatory particle (RP) that caps the CP on one or both ends is responsible for substrate recognition and ATP-dependent substrate unfolding and translocation into the CP[4–6]. Many biological processes are dependent on the proteasome through controlled degradation of key factors, including homeostasis, unfolded protein response and proteostasis[7]. As such, cellular proteasomes are responsible for degrading damaged proteins, thereby preventing the accumulation of misfolded and amyloidogenic proteins, which have a propensity to form aggregates[8].

Aggregation of amyloidogenic proteins progresses through several stages, during which protein monomers assemble into soluble aggregates (oligomers) that through further aggregation events eventually undergo conformational rearrangement into filamentous aggregates (fibrils). The process of protein aggregation is harmful to normal cell physiology and is often associated with neurodegenerative disorders[9]. At the cellular level, accumulation of aggregates could be due to an increased rate of aggregation or decreased rate of aggregate removal, due to e.g. changes in the ability to disassemble or degrade aggregates. Aggregates assembled from amyloidogenic proteins tau and α-synuclein (αS) have been implicated in Alzheimer’s (AD) and Parkinson’s disease (PD), respectively[10,11]. Both tau and αS are intrinsically disordered in their non-amyloid state as monomers, and have been reported to be degradation-resistant as aggregates[12–15].

The inability to process certain aggregates may be coincident with proteasome malfunction, which in certain brain regions of AD and PD patients have been reported with decreased activity[16,17]. We recently demonstrated that the mammalian proteasome holoenzyme possessed a fibril-fragmenting activity, reducing the size of large tau and αS fibrils into smaller entities *in vitro*[1]. Importantly, the proteasome catalyzed this fibril-fragmenting process in a Ub-independent manner. It is currently unclear how these smaller aggregate entities may be further processed by the cellular mechanisms. A recent study has further detailed the interactions of small soluble aggregated amyloidogenic proteins (oligomers) with the proteasome, which is markedly impaired by oligomer binding[18].

Studies in cells have indicated that both tau and αS could be degraded by the proteasome in a Ub-dependent manner[19–22], suggesting that aggregates of ubiquitinated proteins may accumulate when proteasomal functions are compromised. This assumption is supported by the observation of abundantly monoubiquitinated tau fibrils isolated from AD patient brain samples[23]. In addition, αS in the PD-associated Lewy Bodies is also mainly monoubiquitinated[17,24]. Both tau and αS have dedicated Ub ligases, AXOT/MARCH7[25] and SIAH1[26,27], respectively, which preferentially monoubiquitinate their substrates. UBE2W, a Ub-conjugating enzyme that directly monoubiquitinates the N-terminus of intrinsically disordered proteins[28], has also been shown to modify tau[22,23]. Given that proteasomes have affinity for Ub molecules, it is plausible to hypothesize that Ub modification on aggregates would recruit and become targeted by functional proteasomes.

Here we show that the mammalian proteasome holoenzyme can target oligomers assembled from ubiquitinated tau aggregation domain (tau^K18^) and αS. We found that both tau^K18^ and αS may become ubiquitinated on the N-terminus by UBE2W. Using genetically engineered proteins with an N-terminal Ub moiety on tau^K18^ and αS, we demonstrated that such Ub modification delayed the aggregation process, which resulted in distinct aggregate structures compared to their unmodified counterparts. In addition, proteasomal functions were maintained in the presence of these Ub-modified aggregates. This was supported by data from single-molecule fluorescence spectroscopy experiments, which found a reduction in the number and the size of oligomers following proteasome treatment. The ability to target oligomers was not affected by Velcade-mediated inhibition of proteasomal proteolytic activity, suggesting that oligomer disassembly is not dependent on degradation. Based on these observations, we propose that N-terminal Ub modification on tau and αS enables proteasomes to target and remove oligomers assembled from these modified proteins.

## Results

### N-terminal Ub modification on αS and tau^K18^ delays protein aggregation

We chose to use full-length αS and a tetra-repeat domain of tau (tau^K18^) as model amyloidogenic proteins since both protein constructs have a similar molecular weight (∼14 kDa, Figure 1a). Using established protocols to purify untagged recombinant proteins of wild-type αS and tau^K18^, we found that both proteins could be ubiquitinated by UBE2W (Figure 1b). The reaction did not continue beyond monoubiquitination as UBE2W specifically recognizes disordered sequences at the N-terminus of the substrate.

**Figure 1.**
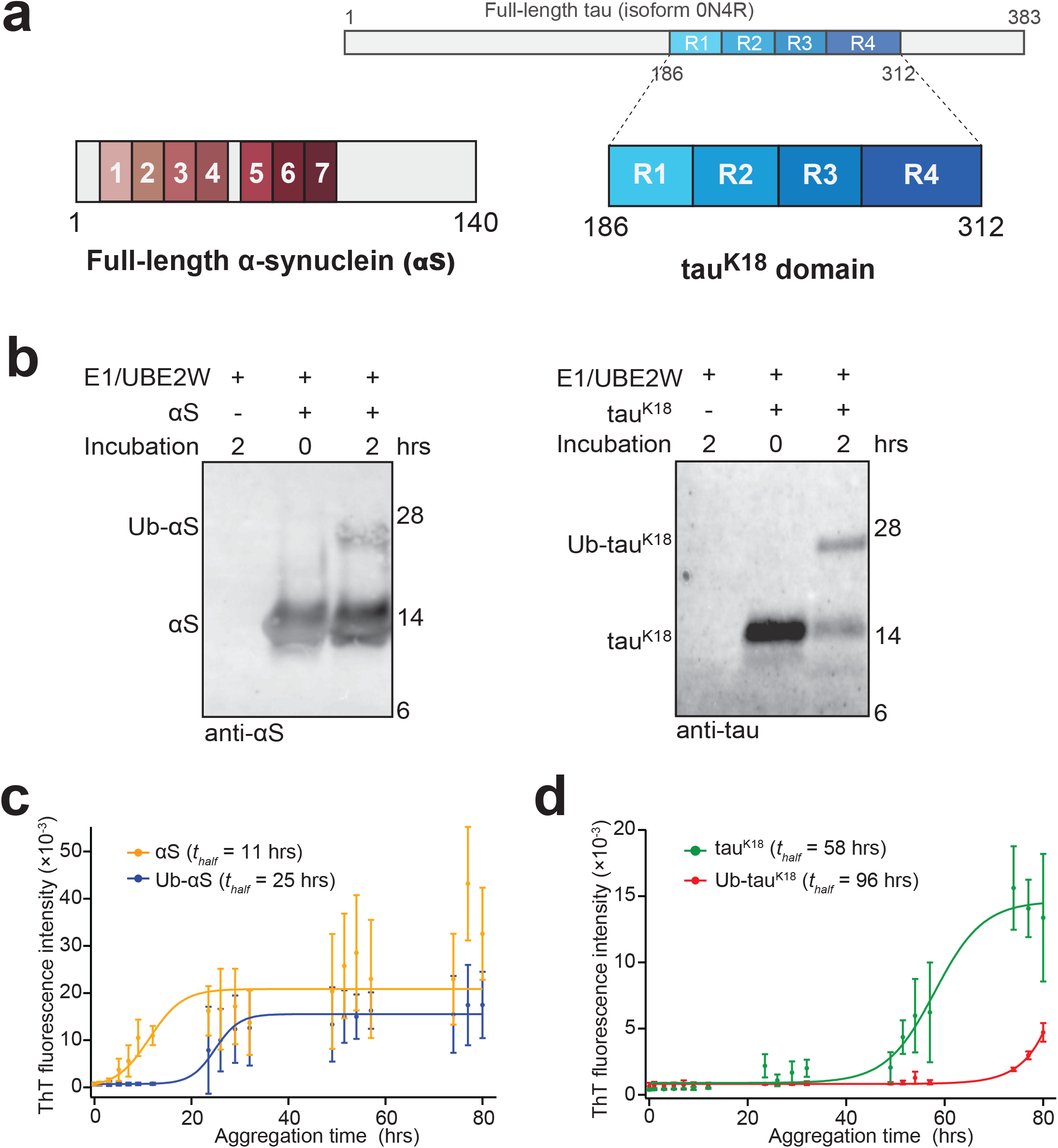
Aggregation of N-terminally Ub-modified tau and α-synuclein (αS). **(a)** Full-length αS (containing at seven repeats) and the tetra-repeat domain (tau^K18^) of tau. Full-length tau (isoform 0N4R) is shown on top. **(b)** Ubiquitination of αS (*left*) and tau^K18^ (*right*) by UBE2W. UBE2W faithfully ubiquitinated tau^K18^ and αS after two hours incubation at 25°C, demonstrated by the shift in the band size. Results shown are representative of reactions independently repeated at least three times. **(c-d)** Aggregation of unmodified and Ub-modified tau^K18^ and αS, detected by Thioflavin T (ThT). **(c)** Ub-αS (blue) or αS (yellow) at 40 µM were aggregated under identical conditions with shaking at 37°C. **(d)** Ub-tau^K18^ (red) or tau^K18^ (green) at 10 µM aggregated under identical conditions, without shaking, at 37°C. Error bars represent standard deviation of independent triplicate measurements.

Protein ubiquitination by UBE2W did not reach completion after 2 hrs, resulting in a two-component mixture of Ub-modified and unmodified protein. To obtain homogenous and pure Ub-modified αS and tau^K18^, we genetically engineered constructs that expressed fusion proteins with a single Ub moiety immediately before the first residue of αS or tau^K18^ (Ub-αS and Ub-tau^K18^, SFigure 1). These engineered N-terminal Ub fusion proteins were protected from deubiquitination by a Gly76Ser substitution of the C-terminal residue of Ub. We further separately cloned the sequences of αS and tau^K18^ alone and purified these unmodified recombinant proteins using the same procedure as for the engineered Ub-modified proteins for consistency (see **Materials and Methods**).

Ub-αS and Ub-tau^K18^ were allowed to aggregate under identical conditions as their unmodified counterparts and measured by ThT, which bound to β-sheet rich amyloid structures. Unmodified αS entered an exponential phase reaching a half-saturated ThT intensity (*t_half_*) at ∼11 hrs (Figure 1c). In contrast, the aggregation of Ub-αS showed an extended lag phase, and reached half-saturation with a delay of 14 hrs (*t_half_* =25 hrs). This Ub-dependent delay was even more apparent for Ub-tau^K18^ (estimated *t_half_* of 96 hrs), whose aggregation was delayed by ∼38 hrs compared to tau^K18^ (*t_half_* =58 hrs, Figure 1d). These results suggest that Ub modification might decrease the rate of aggregate formation and/or the level of total amyloid aggregates under our reaction conditions.

Aggregates assembled beyond 96 hrs were further imaged under TEM to qualitatively compare the effect of Ub modification. Interestingly, while fibrils were detected from unmodified αS, those formed from Ub-αS mostly appeared as small amorphous assemblies (Figure 2a). Despite repeated attempts, we could not detect any filamentous aggregates from Ub-αS under TEM. In comparison, Ub-modified tau^K18^ assembled into aggregates that appeared thinner and less filamentous-like than unmodified tau^K18^, which were detected abundantly (Figure 2b). These results indicate that the morphology of filamentous aggregates is affected by Ub modification.

**Figure 2.**
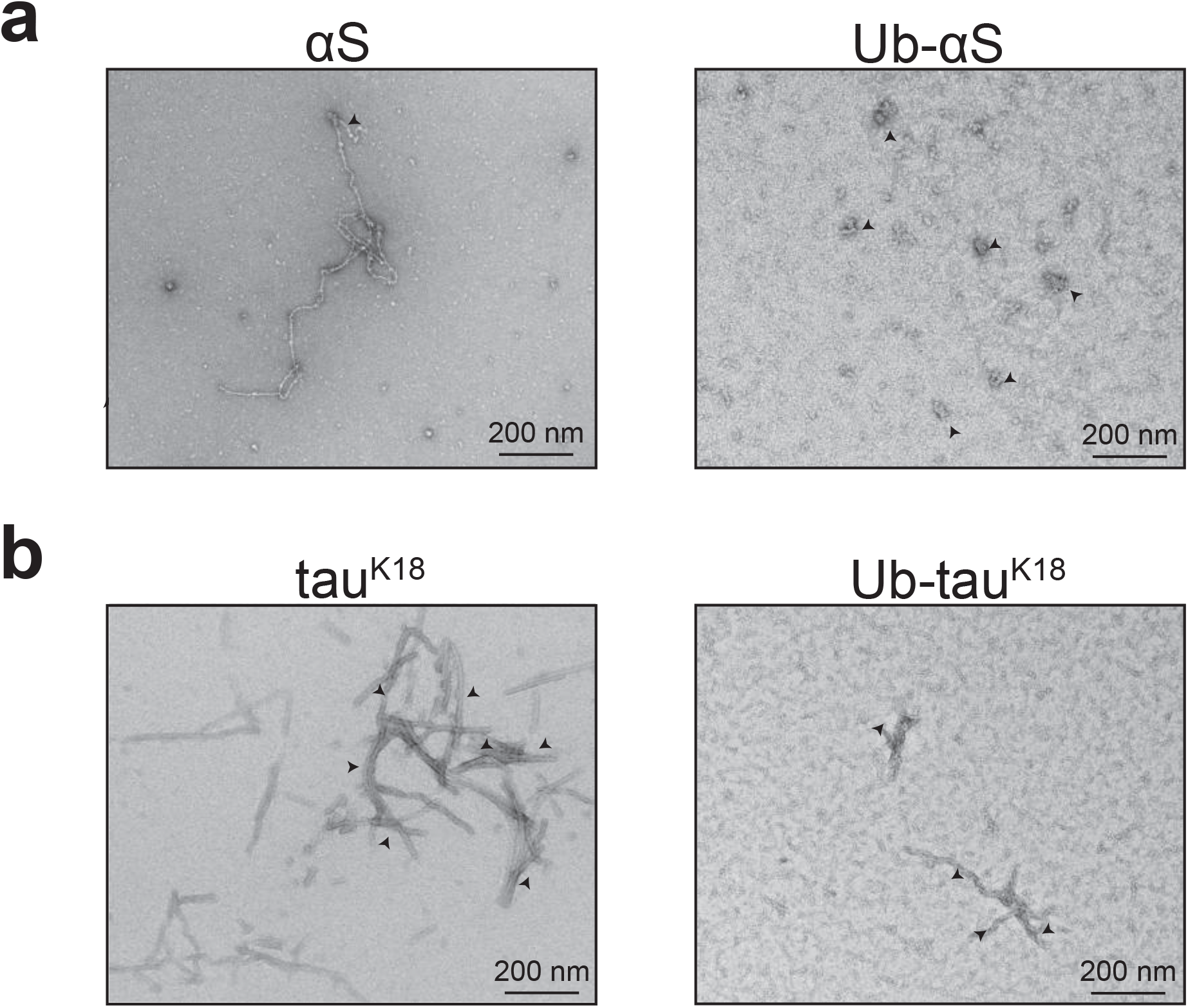
Detection of aggregates by transmission electron microscopy (TEM). **(a)** Filamentous or amorphous aggregates assembled from αS (*left*) and Ub-αS (*right*) after 96 hrs of aggregation reaction. Arrows highlight positions of some typical aggregate structures. **(b)** Filamentous aggregates of tau^K18^ (*left*) and Ub-tau^K18^ (*right*) detected by TEM as in **(a)**.

### Single-molecule measurements of Ub-modified aggregates

We previously established single-molecule fluorescence methods to measure the aggregation process independently of aggregate structure and to estimate the proportion of soluble aggregates (oligomers)[29–31]. In this study, we applied the same method and labeling strategy to attach fluorescent dyes to αS and tau^K18^ (SFigure 2a). Using this approach, we mixed the same protein labeled with Alexa488 or Alexa647 in a 1:1 molar ratio and initiated the aggregation reaction. Aggregate samples were flowed through a microfluidic channel and excited at suitable wavelengths with two overlapping lasers using a confocal microscope (SFigure 2b). Oligomers (here defined as 2-150mers) formed during aggregation will contain both dyes and give rise to coincident fluorescent bursts when they pass through the confocal volume of the laser (Two Color Coincidence Detection (TCCD))[31], while any monomer signal will not give rise to coincident fluorescent bursts (Figure 3a). The fraction of all fluorescence bursts that are coincident is proportional to the fraction of oligomers present and is measured by the *association quotient Q* (see **Materials and Methods**).

**Figure 3.**
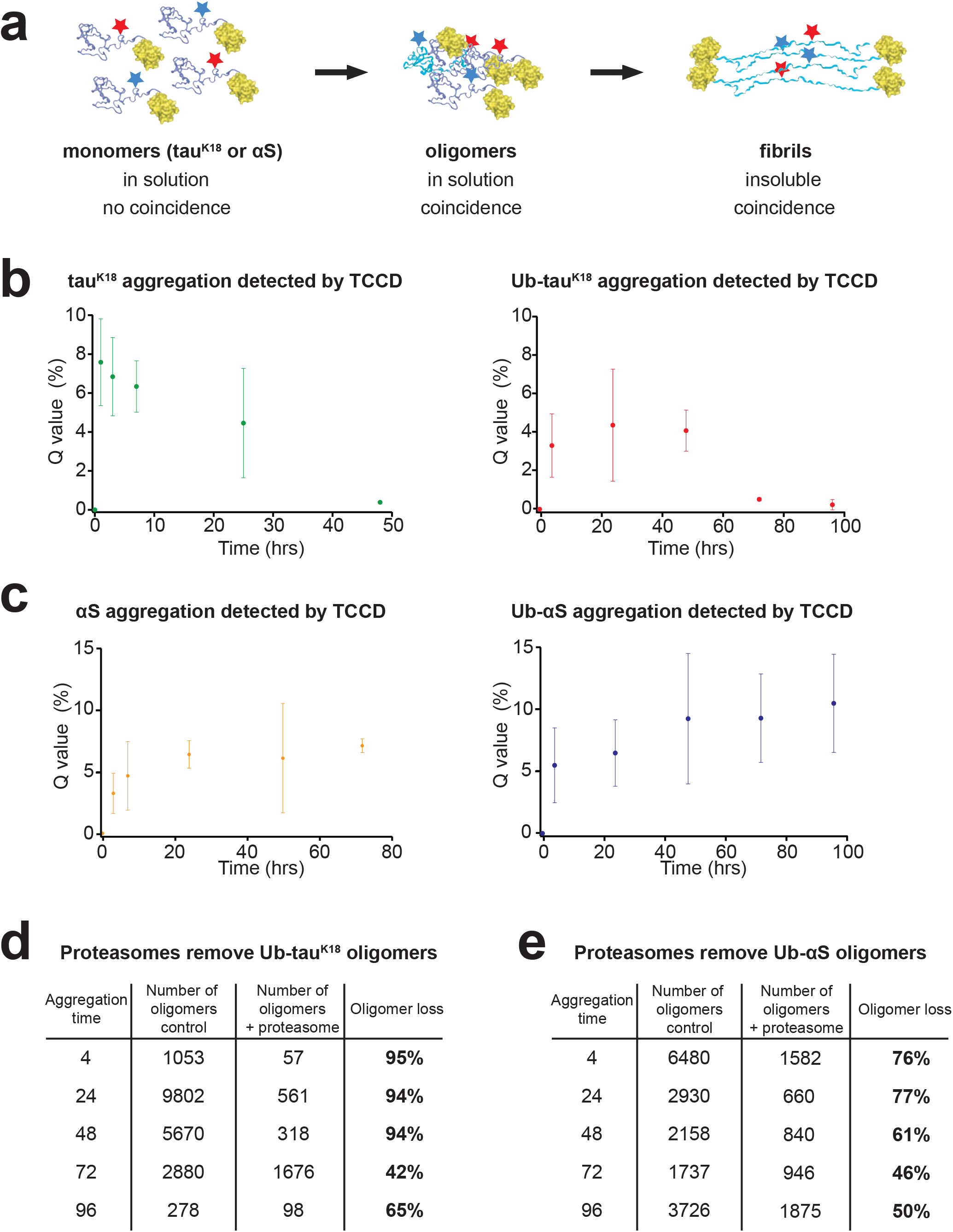
Single-molecule fluorescence detection of αS and tau^K18^ aggregates. **(a)** Schematic representation of aggregation from N-terminally Ub-modified amyloidogenic proteins (in blue; Ub in yellow) tagged with Alexa488 (marine stars) and Alexa647 (red stars) in a 1:1 molar mixture. Monomeric proteins (*left*) carry a single fluorescent dye and cannot be detected using the coincidence criterion **(Materials and Methods)**. Soluble oligomers (*middle*) will carry both dyes and satisfy the coincidence criterion. As aggregation progresses, fibrillar aggregates (in cyan; *right*) will form that may be insoluble or too large for detection. **(b)** Aggregation of tau^K18^ (*left*) or Ub-tau^K18^ (*right*) as detected by single-molecule measurements. The *Q* value is proportional to the percentage of oligomers. Error bars represent standard deviation of three independent measurements. **(c)** Aggregation of αS (*left*) or Ub-αS (*right*) detected by single-molecule measurements as in **(b)**. **(d** and **e)** Representative TCCD experiments of proteasomal degradation of **(d)** Ub-tau^K18^ and **(e)** Ub-αS aggregates, respectively, illustrating how the single-molecule data were analyzed (see **STable 1** for aggregate size dependency analysis). The experiments were performed in triplicate (see Figure 4 for average results of repeat experiments). At the indicated times after aggregation initiation (first column), the number of aggregates were counted after incubation with either the control buffer (second column) or with the proteasome (third column). The calculated percentage reduction of aggregates is shown (fourth column).

We could reproducibly detect oligomers from both tau^K18^ and Ub-tau^K18^ early in the aggregation process (Figure 3b). Oligomers assembled from tau^K18^ reached an estimated peak level between ∼1-3 hrs after reaction initiation. This observation is qualitatively consistent with our previously published results[30]. For Ub-tau^K18^ the highest oligomer level was detected between ∼24-48 hrs of incubation. An apparent reduction of the fraction of oligomers in solution with time was detected in both unmodified and Ub-modified tau^K18^. The loss of soluble aggregates as aggregation progresses had previously also been observed for tau^K18^ without Ub modification[30], and could be due to presence of aggregates that are either insoluble or too large to enter the microfluidic channel and hence not detected in our single-molecule experiments.

A steady state population of soluble oligomers appeared in the aggregation of both unmodified and Ub-modified αS and did not change appreciably as the reaction proceeded over longer incubation times (Figure 3c). Only a fraction of the aggregates is therefore able to form ThT-active aggregate species, where the formation of Ub-modified αS aggregates is delayed compared to unmodified αS (shown in Figure 1c). The ThT assays further suggest that there could be more β-sheet content in the aggregates formed from unmodified αS, as a bulky Ub moiety in Ub-αS may obstruct close packing.

### Proteasomes are able to target Ub-modified aggregates

To study the degradation of Ub-modified amyloidogenic proteins, we purified proteasome holoenzyme from an established mammalian cell line[32] using an affinity column (SFigure 3a). The purified proteasome holoenzyme was resolved and validated by SDS-PAGE (SFigure 3b), transmission electron microscopy (TEM, SFigure 3c) and native gel electrophoresis in presence of an ATP-containing buffer (SFigure 3d). We could not quantitatively detect the presence of free CP with Coomassie staining or under TEM. A batch of the yeast proteasome holoenzyme was used as reference for the detection of capped and uncapped CP under identical conditions (SFigure 3e). All four Dye-labeled tau^K18^ and αS protein constructs could be quantitatively degraded by the proteasome, its activity (SFigure 4a and b).

**Figure 4.**
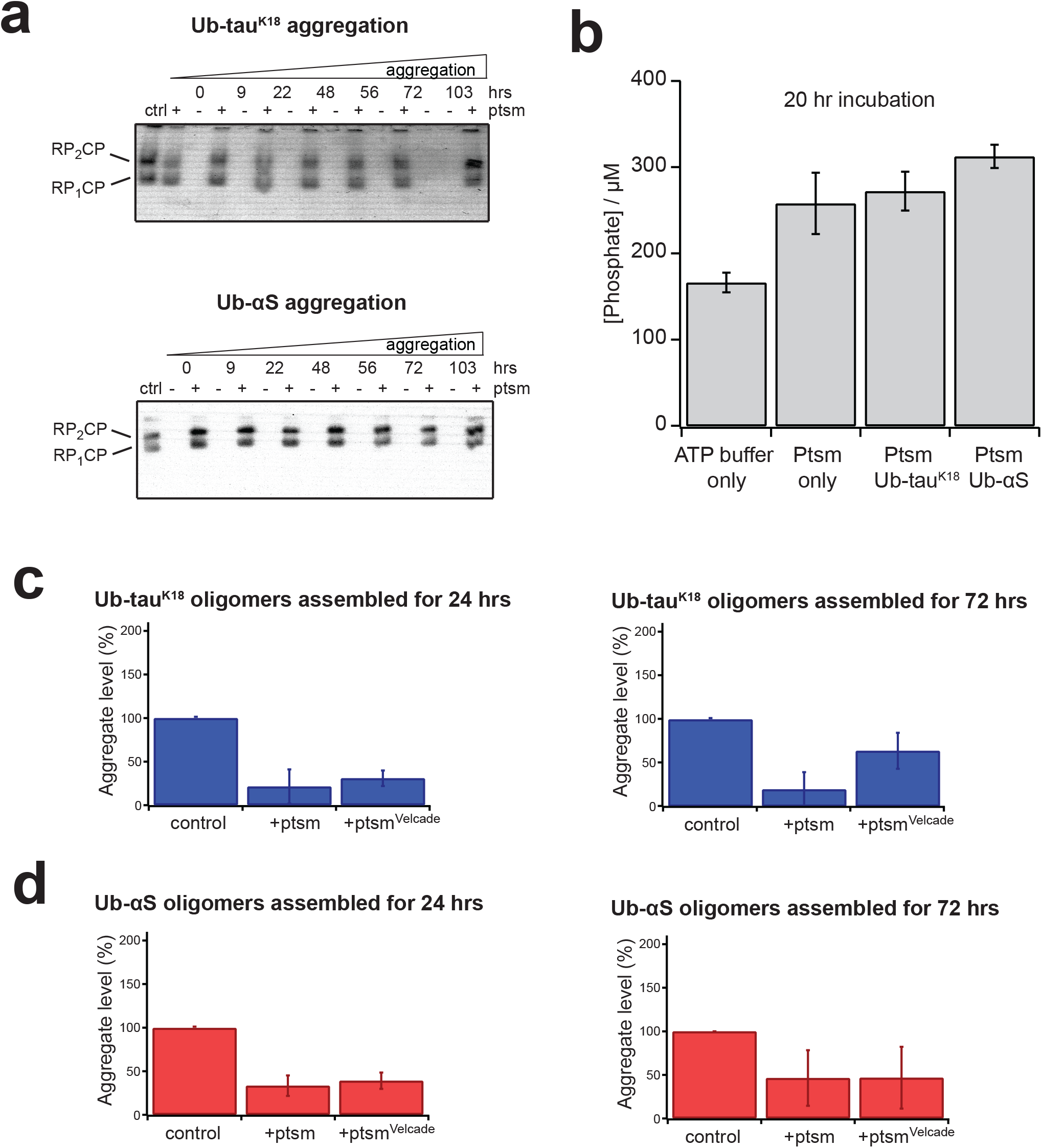
Proteasome maintains its function in the presence of Ub-modified oligomers, which are disassembled independently of the proteolytic acitivity. **(a)** Ub-tau^K18^ (*top*) or Ub-αS (*bottom*) aliquots from indicated aggregation times were incubated with the proteasome or with control buffer. After reaction completion, samples were resolved on a 3% native gel and visualized by LLVY-AMC fluorescence emission (_λ*Ex*_ = 340 nm, _λ*Em*_ = 440 nm). A sample of the proteasome at the same concentration alone was used as control (ctrl). **(b)** Phosphate assay reporting the concentration of free phosphates after incubating proteasomes with Ub-modified oligomers for 20 hrs. Residual free phosphates were present in the ATP-containing buffer, but increased in presence of the proteasome. The free phosphate levels were not reduced when the proteasome was incubated with Ub-modified tau^K18^ or αS, suggesting that ATP hydrolysis was not affected. **(c-d)** Velcade does not affect aggregate removal by the proteasome. **(c)** Ub-tau^K18^ and **(d)** Ub-αS oligomers were assembled for 24 hrs (*left*) or 72 hrs (*right*) before mixing with the prepared proteasomes, incubated and subsequently measured by single-molecule TCCD. Proteasomes were pre-incubated with Velcade for 5 min before mixing with substrates. The percentage change in aggregate level is relative to the control sample without proteasome. Error bars represent standard error of mean from three sets of independent measurements.

It has been reported that non-ubiquitinated aggregates resist degradation[12–15], and that tau and αS oligomers have recently been shown to impede proteasome activity[18]. In addition, we recently demonstrated that the product of proteasome-catalyzed fragmenting of tau and αS fibrils was small aggregate entities, supporting our previous report that proteasomes had no effect on soluble αS oligomers without ubiquitin modification[39]. We therefore tested whether N-terminal Ub modification on tau and αS oligomers would enable their disassembly by proteasomes. ThT was found to bind to the proteasome, which interfered with fluorescence measurements (SFigure 4c). Turning to single-molecule TCCD approach, we reproducibly detected a decrease in the level of soluble oligomers in both Ub-tau^K18^ and Ub-αS as late as 96 hrs into the aggregation reaction (a typical set of experimental data is shown in Figure 3d and e). Ub-tau^K18^ oligomers generated throughout the assembly process were largely removed after incubation in the presence of proteasomes (Figure 3d). The effect was also significant for oligomers assembled from Ub-αS, which decreased in the presence of the proteasome (Figure 3e).

We previously used the relative fluorescence intensities of aggregates compared to monomers to calculate the apparent size of aggregates[29]. Analyzing the data in Figure 3c and d revealed that the proteasome caused a general decrease in the aggregate level independently of the apparent aggregate size (**STable 1**), suggestive of gradual aggregation disassembly and reduction.

Our previous work using the same confocal single-molecule technique found that the proteasome did not target oligomers that are not modified with Ub[12] and that these oligomers are not affected by active chaperones[33]. We therefore attribute the observed reduction in aggregate level specifically to the proteasome. No change in the proteasomal proteolytic or the ATP-dependent activity after incubation with Ub-modified oligomers was observed (Figure 4a and b), consistent with observations after incubating proteasomes with fibrils[1].

The fibril-fragmenting activity of proteasome was shown to be independent of the peptidase activity[1]. To test whether the disassembly of Ub-modified aggregates also did not rely on the proteasomal peptidase activity, we repeated the single-molecule experiments using proteasomes that were pre-incubated with Velcade as before. The decrease in Ub-tau^K18^ or Ub-αS aggregates was not affected by Velcade-mediated inhibition (Figure 4c and d), consistent with our recent observation[1].

The ability of proteasomes to target Ub-modified oligomers was further resolved by Western blotting. Aggregates assembled from both Ub-tau^K18^ and Ub-αS were prone to proteasomal activity (SFigure 5a and c), while aggregates assembled from unmodified tau^K18^ and αS showed no detectable intensity change in the presence or absence of proteasome treatment (SFigure 5b and d). This is consistent with the hypothesis that aggregates formed from proteins N-terminally modified with Ub can be targeted by the proteasome while unmodified aggregates remain unchanged.

## Discussion

Our current work has showed that the presence of an N-terminal Ub moiety on tau^K18^ and αS results in distinct aggregation kinetics and aggregate conformations. The oligomers assembled from these Ub-modified proteins are also prone to proteasomal activity, leading to reduction in their levels. We previously demonstrated that proteasomes are able to target fibrils, as they are structurally distinct from non-filamentous aggregates[1]. This present study further complements our previous results with an additional proteasomal ability to target Ub-modified oligomers.

As tight assembly and packing are key features of amyloid aggregates, presence of a Ub moiety at the N-terminus could potentially induce soluble oligomers into a conformation distinct from unmodified protein aggregates. In another of our previous work, we found that arachidonic acid could induce a conformational change in soluble αS oligomers that could subsequently be targeted by the proteasome[12], suggesting that the proteasome acted more effectively on modified than unmodified oligomers, which contained more compact structures.

In addition to restricting or delaying the formation of proteasome-resistant aggregate conformations, it is plausible that the presence of Ub moieties would provide affinity for proteasomes to engage oligomers more faithfully. Avid recognition of dispersed Ub moieties on aggregates may contribute to preferential recognition followed by disassembly of Ub-modified over unmodified aggregates. Multiple Ub ligases have been proposed to monoubiquitinate amyloidogenic proteins, including UBE2W-mediated N-terminal modification[28,34], which may serve as a mechanism to counteract protein aggregation and associated cell damage. Our current study provides a possible option for how oligomers may become disassembled by the mechanisms of the ubiquitin-proteasome system.

## Supporting information

SFigures 1-5; STable 1

## Acknowledgements

The authors would like to thank members of the D.K. and D.F. labs for reagents and helpful discussions. Y.Y. acknowledges a Henslow Research Fellowship from Selwyn College, Cambridge. This research is funded by a Sir Henry Wellcome Research Fellowship awarded to Y.Y. Research carried out in DF’s lab is funded by NIH grant R01 GM043601 and a grant from the Rainwater Foundation.

## Declarations

The authors declare that they have no conflict of interest.

## Author Contributions

D.F. and D.K. directed the research. D.F., D.K. and Y.Y. designed the experiments. Y.Y. conceptualized the project, performed the experiments, analyzed the data and prepared the manuscript. All authors contributed to the writing of the manuscript.

---

**SFigure 1.**
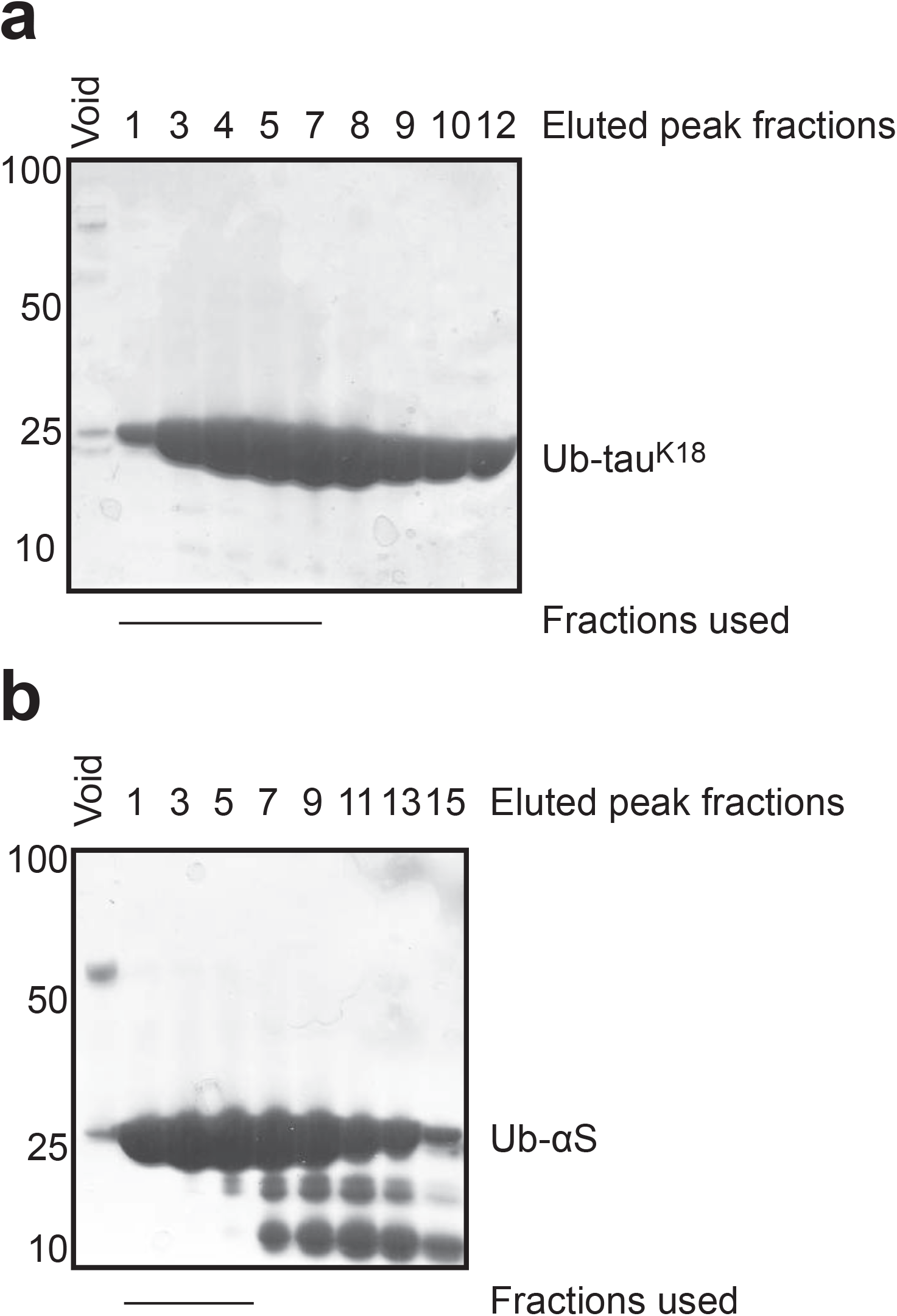
Purification of Ub-tau^K18^ and Ub-αS. **(a)** Peak fractions of Ub-tau^K18^ eluted from a typical run on gel filtration column were resolved on SDS-PAGE and proteins were stained with Coomassie blue. **(b)** Ub-αS samples were also purified and eluted from gel filtration column as in **(a)**. The fractions collected and used for experiments are underlined. Molecular weight markers are shown to the left of each gel.

**SFigure 2.**
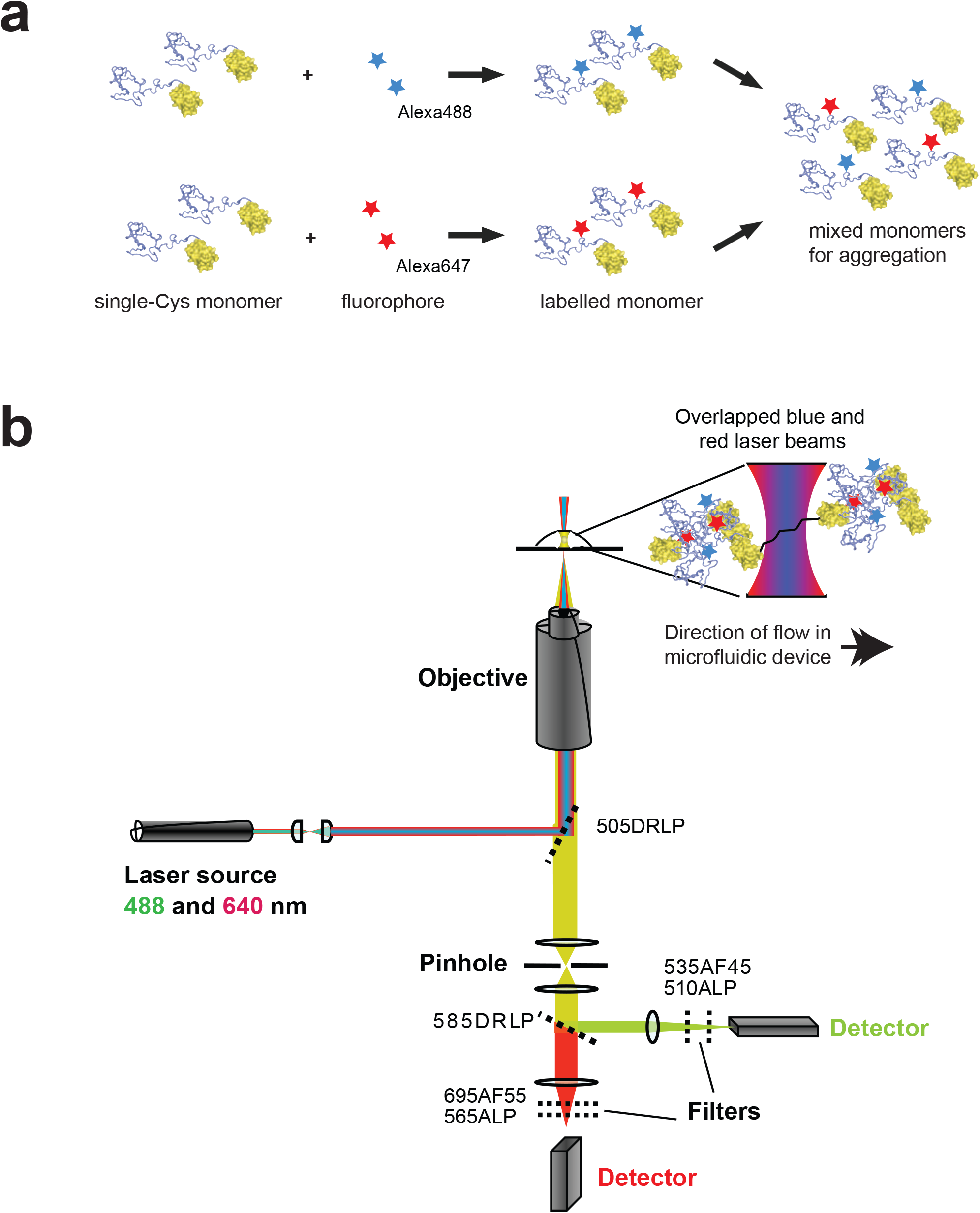
Single-molecule confocal measurements of oligomers. **(a)** Dye-labeling scheme used for Ub-tau^K18^ and Ub-αS. The proteins were genetically engineered to carry a single Cys residue that enables covalent fluorescent dye labelling (see **Materials and Methods**). Each protein was labeled separately with Alexa488 or Alexa647 before mixing in a 1:1 molar ratio for subsequent aggregation. **(b)** Model representation of the single-molecule confocal instrument setup used. A multi-line laser source (iFLEX-Viper) with a modular single mode fibre delivery system was responsible for excitations at 488 nm and 640 nm. The two overlapping laser beams were focused using an objective with high numerical aperture into a microfluidic channel where a constant flow was applied to the protein sample (*zoomed in*). The sample was diluted to a concentration such that single molecules occupying the focal volume were simultaneously excited. Emitted fluorescence is captured using the same objective and separated from reflected excitation beam using a dichroic mirror (DRLP, dichroic long-pass filter). A second dichroic mirror splits the emission light beam again for detection in two separate APDs (avalanche photodiode detector). ALP (anti-light pollution) filters and AF (band pass) filters used are indicated.

**SFigure 3.**
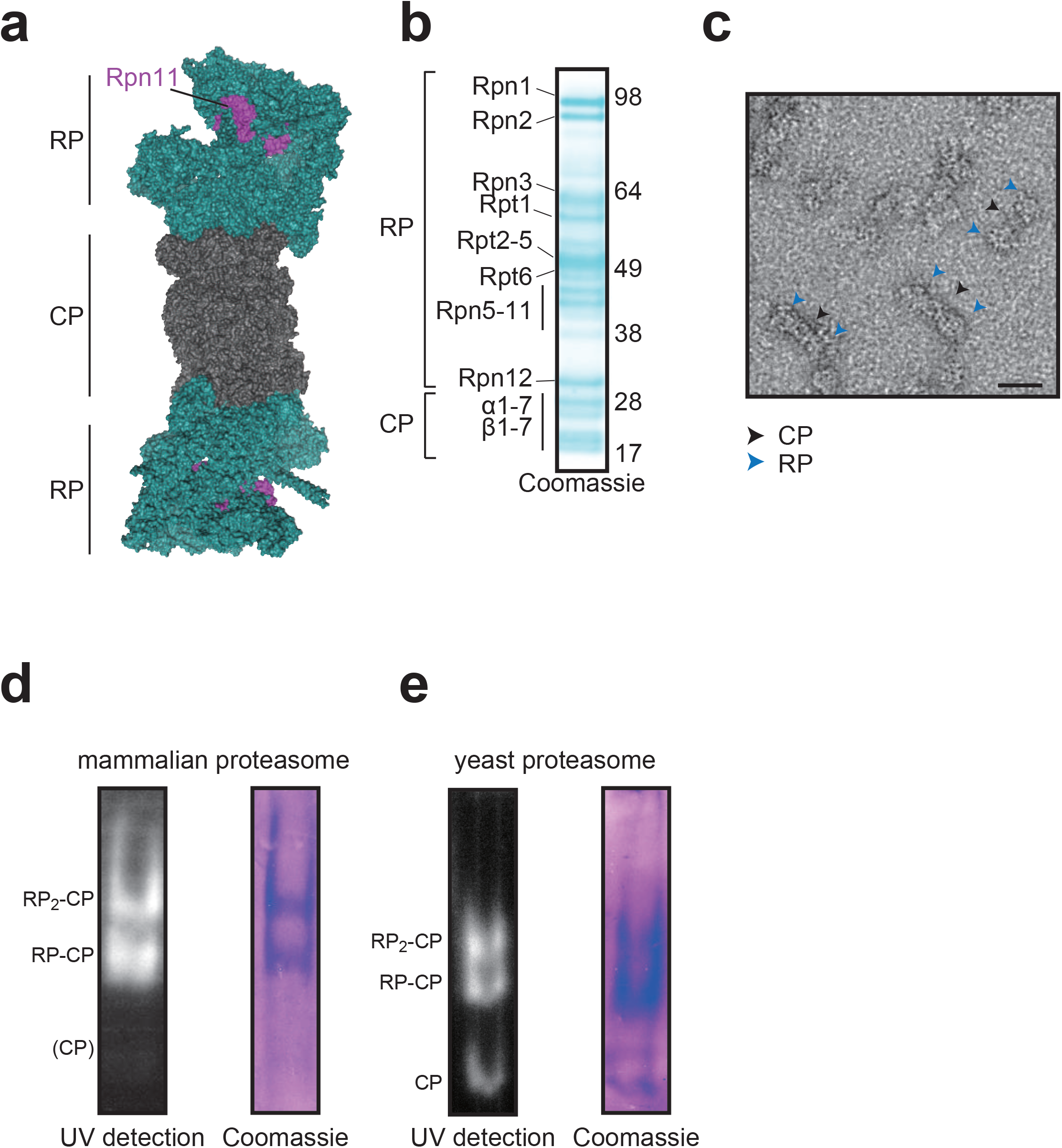
Purification of 26S proteasome holoenzyme from a HEK293-derived cell line. **(a)** Surface representation of proteasome holoenzyme (pdb-id 5GJQ). The 20S core particle (CP, grey) is capped at both ends with 19S regulatory particles (RP, teal). Rpn11 (magenta), a deubiquitinase, is the subunit tagged for affinity purification of the proteasome (see **Materials and Methods**). **(b)** The presence of the individual proteasomal subunits was detected by SDS-PAGE after purification. **(c)** A typical batch of proteasome holoenzyme visualized by TEM. The CP and RPs are highlighted. **(d)** Purified proteasome holoenzymes used in this study resolved by native gel electrophoresis and detected under UV with LLVY-AMC (*left*) or Coomassie-stained (*right*) **(e)** A batch of yeast 26S proteasomes detected under UV (*left*) or stained with Coomassie (*right*).

**SFigure 4.**
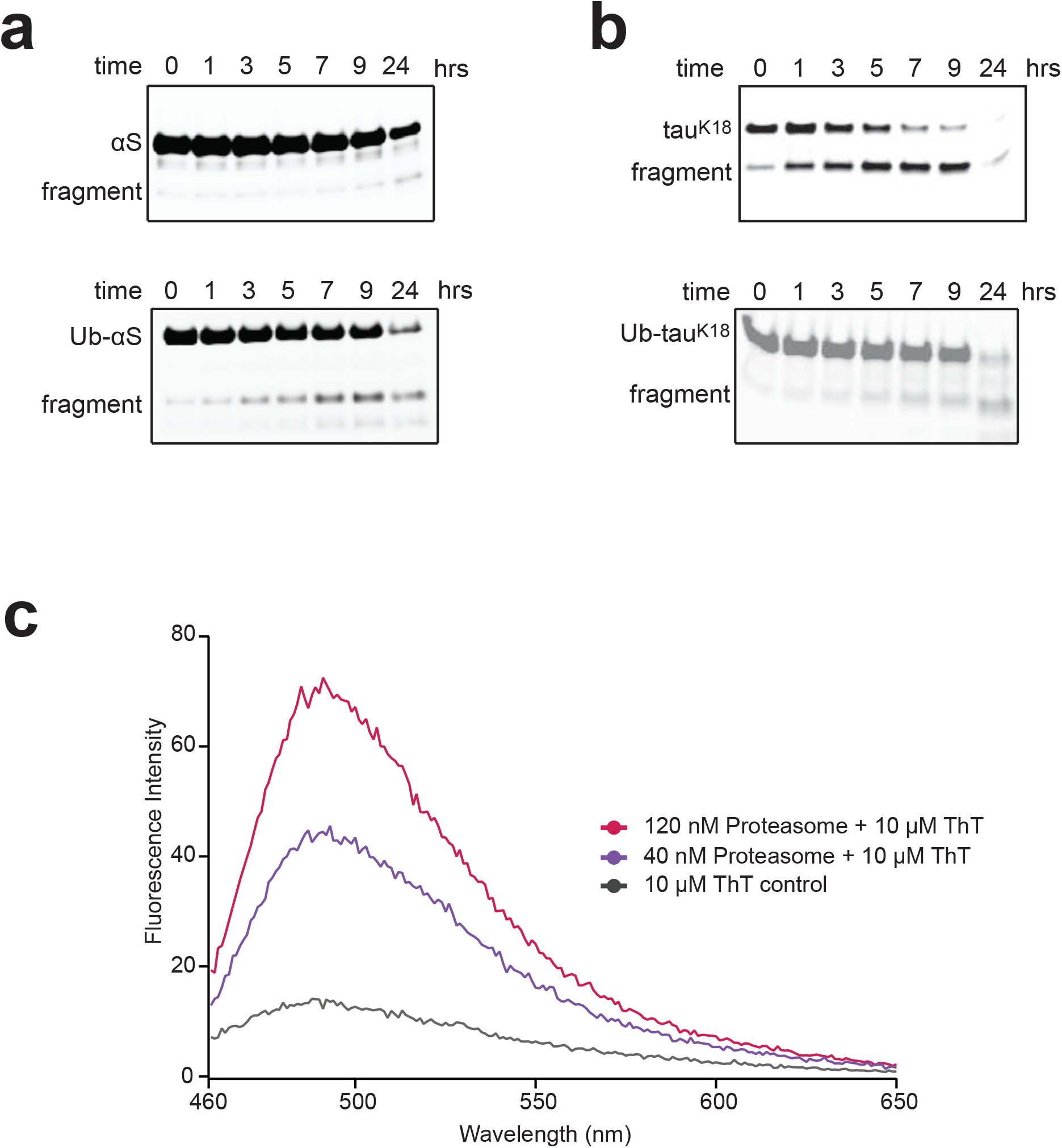
Studying proteasome activities on N-terminally Ub-modified proteins. **(a)** Dye-labeled αS (top) and Ub-αS (bottom) monomers can be degraded by the proteasome over time. Samples were resolved by SDS-PAGE and fluorescence emission was detected on a Typhoon Imager. A lower molecular weight degradation fragment developed over time beneath the substrate bands. (**b**) Degradation of dye-labeled tau^K18^ (*top*) and Ub-tau^K18^ (*bottom*) monomers, performed and visualized as in **(a)**. **(c)** ThT can bind to the proteasome. The proteasome incubated at 0, 40, or 120 nM with ThT at 10 µM final concentration. Samples were excited at 415 nm and fluorescence emission was detected from 460 to 560 nm.

**SFigure 5.**
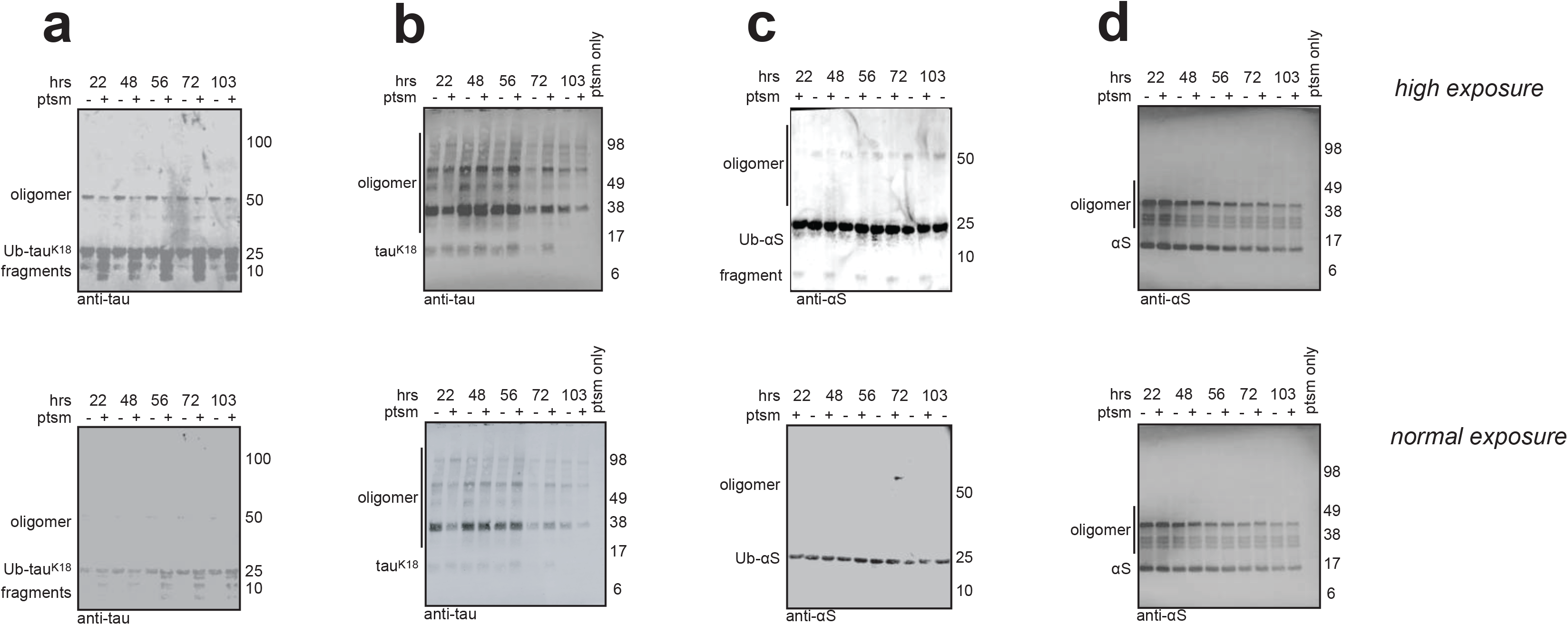
Degradation assays of Ub-modified oligomers. Degraded samples were resolved by SDS-PAGE. Oligomers assembled from **(a)** Ub-tau^K18^ or **(b)** tau^K18^ were incubated with the proteasome or the control buffer, resolved by SDS-PAGE and transferred to a PVDF membrane for Western blot detection and high (*top*) and normal exposure (*bottom*) to detect changes in the oligomer or monomer bands. Oligomers assembled from **(c)** Ub-αS or **(d)** αS were subjected to proteasomal degradation and presented as in **(a** and **b)**. Note that the proteasome alone did not contribute to any bands detected by either anti-tau or anti-αS antibodies. The reduction in oligomer bands of Ub-tau^K18^ and Ub-αS aggregates throughout the aggregation process was not an effect of the reducing agent or other factors because samples in control lanes and proteasome-containing lanes underwent the same procedure and were kept in the same buffer. Each assay was independently repeated at least three times.

**Supplementary Table 1.**
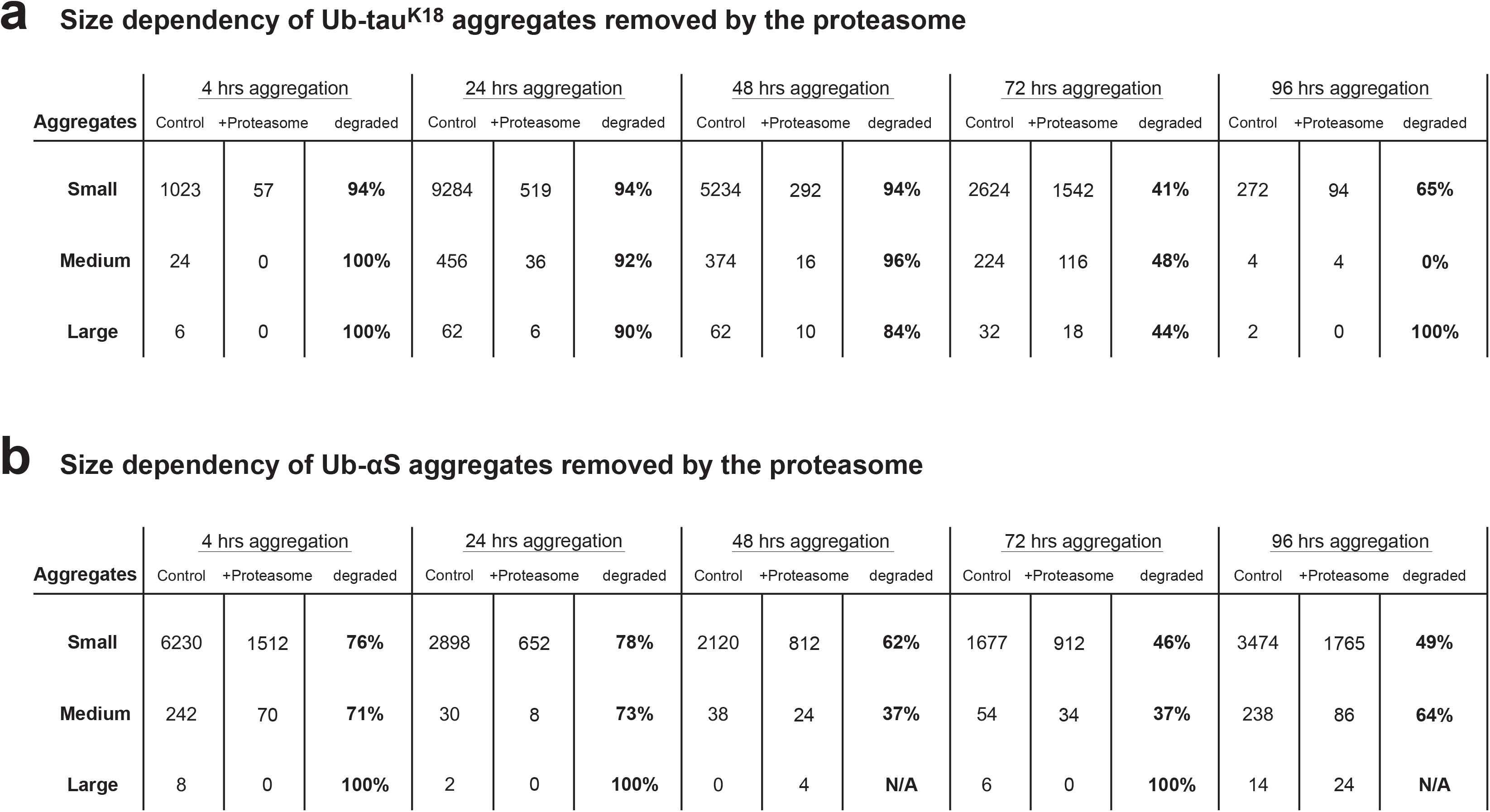
Changes in oligomer size distribution following proteasomal degradation calculated from data in Figure 3d and e. **(a)** Ub-tau^K18^ and **(b)** Ub-synuclein oligomers are grouped according to their *apparent oligomer sizes* into ‘*Small’* (15mers or smaller), ‘*Medium’* (between 15mers and 30mers), and ‘*Large’* (between 30mers and 45mers).

## Material and Methods

### Molecular biology of plasmids for protein expression

A single Ub moiety was expressed in tandem to the proteins of interest (tau^K18^, αS or full-length tau). The DNA sequence of Ub was introduced either at the 5’ end, immediately before the ATG codon, or at the 3’ end, after the last translated codon of the open reading frame. When the Ub coding sequence was cloned upstream of the wild-type αS or tau^K18^ coding sequences, a mutation corresponding to Gly to Ser was introduced at residue 76, the last residue of Ub. The constructs were subsequently sub-cloned into a pOPINF vector using restriction enzymes, resulting in a His_6_-tag at the N-terminus of the Ub. Tau^K18^ or αS sequences were also separately cloned into the pOPINF vector without the Ub sequence, so that the His_6_-tag is immediately N-terminal to the substrate.

Cys mutations were introduced using site-directed mutagenesis on Ala90 of αS or Ile202 of tau (annotation based on the 0N4R isoform sequence). We previously showed that introduction of Cys and subsequent dye-labeling did not disrupt the integrity and aggregation properties of these proteins[29,30]. Equivalent mutations were separately introduced into Ub-αS and Ub-tau^K18^ constructs. The two other Cys residues in wild-type tau^K18^ sequence were mutated to Ala[30].

The full-length sequence of mouse E1 in pET28a vector (kind gift from David Komander) and human UBE2W in pET15b vector (kind gift from Wade Harper, Addgene plasmid #15809) were coded to include an N-terminal His_6_-tag to facilitate protein expression. Plasmids for protein expression of full-length αS in pT7-7 vector or tau^K18^ sequence in pJExpress vector (custom designed) alone coded for untagged constructs of the wild-type sequence of human αS or tau^K18^. These untagged constructs coded for wild-type sequences of the N-terminal residues used in the ubiquitination by UBE2W.

### Recombinant protein purification

Plasmids were transformed into Rosetta2 (DE3) pLysS cells (Novagen) and grown in LB media to OD_600_ = 1.0 before overnight induction with 1 mM IPTG at 20°C. Cells were collected the next day by centrifugation at 5000 × *g* for 30 min before lysis by sonication. The cell lysate was cleared by centrifugation at 21000 × *g* for 30 min at 4°C.

The supernatant from purification of His_6_-tagged proteins was loaded onto a self-packed cobalt column (Clontech). Unbound proteins were washed off with Loading Buffer (50 mM Tris-HCl [pH 7.4], 100 mM NaCl, 10 mM imidazole), and bound proteins subsequently eluted with Elution Buffer (50 mM Tris-HCl, 100 mM NaCl, 200 mM imidazole, pH adjusted to 7.4).

Preparation of the supernatant from purification of untagged αS and tau^K18^ followed established protocols (e.g. [29,35]). In brief, the cleared supernatant was poured into a 50 ml falcon tube and incubated in boiling water for 10 min before cooling down to room temperature. The solution was cleared with a second centrifugation step at 21000 × *g* for 30 min at 4°C and the supernatant filtered before further purification.

The eluted or filtered samples were further purified using ion exchange (IEX) chromatography columns HiTrapQ (for αS constructs) or HiTrapS (for tau constructs), running a linear NaCl gradient from Buffer A (50 mM Tris-HCl [pH 7.4], 50 mM NaCl) up to 1 M NaCl.

Peak fractions from the IEX were concentrated to < 4 ml and loaded onto a Superdex 16/60 gel filtration column (GE Healthcare) in Buffer A. Eluted fractions were separated by SDS-PAGE and fractions judged to be >99% pure by Coomassie stain were further concentrated and flash frozen in aliquots (SFigure 1). Protein concentrations were measured on a NanoDrop.

### Dye-labeling on proteins

Proteins carrying Cys substitutions were dialyzed into Labeling Buffer (50 mM Tris-HCl [pH 7.2]) before dye labeling. AlexaFluor 488 C5 maleimide or AlexaFluor 647 C2 maleimide (Invitrogen) were dissolved in DMSO and added to the proteins in a 1:1.2 molar ratio of excess dye. We routinely use this protocol to ensure that essentially all proteins are labeled as detected by ion exchange chromatography, size exclusion chromatography, and mass spectroscopy. The labeling reaction was quenched after 1 hr with fresh DTT at 100 mM final concentration and loaded onto a HiPrep 26/10 desalting column (GE Healthcare). The final labeled proteins were concentrated to at least 80 µM and flash-frozen in Protein Buffer (50 mM Tris-HCl [pH 7.2], 50 mM NaCl, 0.01% Tween20) in small aliquots. Concentrations were determined by NanoDrop.

### Purification of mammalian proteasomes

Proteasomes were purified from a HEK293T cell line stably expressing a Rpn11-TEV-Biotin tag (kind gift of Lan Huang, UC Irvine) using established protocols[32]. Briefly, cells were grown to 100% confluence and collected with a scraper before resuspension in ice-cold Proteasome Buffer (50 mM Tris [pH 7.5], 0.5% NP-40, 10% glycerol, 5 mM ATP, 1 mM DTT, 5 mM MgCl_2_,). A Dounce homogenizer was then used to lyse the cells and the lysate was cleared by centrifugation at 3000 × *g* for 5 min at 4°C. The lysate was incubated overnight at 4°C in 2-ml bed volume of pre-equilibrated NeutrAvidin resin beads (Pierce). Unbound proteins were washed off with Proteasome Buffer. Proteasome was subsequently cleaved on the column with TEV protease (Invitrogen) at 30°C and concentrated to > 2 µM before flash-freezing.

### UBE2W ubiquitination assays

Ubiquitination assays were carried out at 25°C in Ubiquitination Buffer (50 mM Tris [pH 7.5], 10 mM ATP, 1 mM DTT, 10 mM MgCl_2_,), containing 0.5 µM of E1, 1 µM of UBE2W, 200 of µM wild-type Ub (Sigma) and 10 µM of untagged αS or tau^K18^ substrate. The reactions were incubated for 2 hrs before being quenched by Laemmli Buffer. Substrates were excluded in control samples to test for cross-reactivity of the antibodies used with the ubiquitinating enzymes or the Ub (Figure 1b).

### Protein aggregation assays

For protein aggregation, Ub-tau^K18^ or tau^K18^ were diluted in PBS Buffer (MP Biomedicals) containing 0.01% sodium azide to 10 µM final concentration and incubated at 37°C. An equimolar amount of heparin (H3393, Sigma-Aldrich) was added to initiate tau^K18^ aggregation reactions. Aggregation assays for Ub-αS or αS were performed at 40 µM final protein concentration in PBS Buffer containing 0.01% sodium azide and incubated at 37°C shaking, as described[29]. We did not detect pellets of insoluble fibrils after 10 min centrifugation at 13000 × *g* for Ub-tau^K18^ or Ub-αS aggregates.

### Thioflavin T fluorescence assays

Thioflavin T (Sigma) was dissolved in PBS and filtered through a 0.02 µm filter. The concentration was determined by UV absorbance at 405 nm on a NanoDrop. Aliquots were removed from tau^K18^ (10 µl) or αS (5 µl) samples at indicated times after aggregation initiation and mixed with 40 µl of the ThT solution at 10 µM. The mixture was incubated for 10 min and subsequently measured on a spectrophotometer (λ_*Ex*_ = 415 nm, Varian Eclipse). Integral area between 460 and 560 nm of the emission spectrum was calculated for each time point. The mean value of triplicate aggregation assays was used to plot Figure 1c and d. Each dataset was fitted to a sigmoidal function, defined as

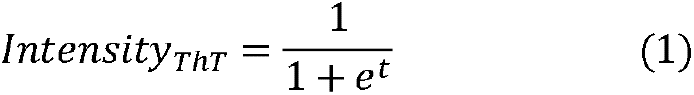

where *t* is the time after aggregation initiation in hrs and *Intensity_ThT_* is the mean integral area of fluorescence emission. All plots were calculated and generated using IgorPro (Wavemetrics). The sigmoidal behavior of aggregation produces the time, *t_half_*, needed for *Intensity_ThT_* to reach 50% of the maximal plateau value.

### Degradation assays

Degradation assays were typically performed at 25°C in Degradation Buffer (50 mM Tris-HCl [pH 7.5], 10 mM ATP-MgCl_2_, 30 mM creatine phosphate, 4 µM creatine kinase) containing 40 nM proteasome. Control samples were set up in the same buffer without adding the proteasome. Tau^K18^ substrates were reacted for 3 hrs at a final concentration of 2.6 µM in PAGE-based assays and 0.2 µM in single-molecule assays. Incubation time for αS samples was 12 hrs and diluted to a final concentration of 10 µM in PAGE-based assays and 1 µM in single-molecule assays. We occasionally observe secondary dimer or higher bands emerging with prolonged incubation when the substrate concentration is higher than 10 µM, indicating sporadic aggregation during degradation assays. For the inhibitor assays, the proteasome was pre-incubated for 5 min at 25°C with Velcade (Proteasome^Velcade^) to a final concentration of 100 µM.

### Resolving of aggregate samples by gel electrophoresis and Western blotting

Proteasomes used in the overnight degradation assays were resolved on self-poured 3% polyacrylamide native gels and detected with a fluorogenic model substrate, LLVY-AMC, as described previously[36]. For Western blotting, protein samples were first separated on 4-12% NuPAGE gels (Invitrogen) and then transferred to PVDF membranes as per manufacturer’s protocol (Mini Trans-Blot wet transfer, Biorad). Samples taken were quenched with Lammeli buffer but not heated (to preserve the oligomeric bands). Mouse anti-tau (1E1/A6, Merck) or rabbit anti-αS (ab138501, Abcam) were used as primary antibodies following standard Western blotting methods. Secondary anti-mouse and anti-rabbit antibodies compatible with detection on an Odyssey CLx Imager or a Typhoon scanner were purchased from Li-Cor or Invitrogen.

### Colorimetric phosphate assay

ATPase kit containing malachite green and ammonium molybdate was purchased from Abcam (ab65622). For assays in Figure 4b, 40 µl of each reaction (set up as described in the section **Degradation assays**) was mixed with 6 µl of the malachite green reagent and incubated for 15 min before measurement. The colorimetric output was measured at OD = 650 nm on a microplate reader. A linear standard curve from 0 to 20 mM of free phosphates was established to convert colorimetric reading into phosphate concentration. Three independent replicate experiments were performed for each reaction.

### Transmission electron microscopy imaging

Samples shown in Figure 2 were aggregated beyond 96 hrs and applied onto a carbon-coated 400 mesh copper grid (Agar Scientific). Mammalian proteasome was applied at 100 nM concentration. The grids were then washed with double distilled water and stained with 2% (w/v) uranyl acetate for 1 min. TEM images were acquired using Tecnai G2 microscope (13218, EDAX, AMETEK) operating at an excitation voltage of 200 kV.

### Single-molecule measurements

The instrument setup and collection of single-molecule data are based on our earlier works and have been extensively described (e.g. refs [29,37]). Briefly, single-molecule data were collected on a custom-built system using overlapping lasers with excitation maxima at 485 nm and 640 nm (see SFigure 2a). The rate of modulation for both lasers was at 10 modulations per millisecond. Samples were measured under flow using custom-made PDMS microfluidics devices following published procedures[38]. Aggregates were assembled from protein monomers labeled with either AlexaFluor 488 or AlexaFluor 647 and mixed in a 1:1 molar ratio for aggregation (SFigure 2a). This ensures that only aggregates will carry both dyes and will be detected by the coincidence criterion[31].

Degradation assays were performed for 3 hrs (Ub-tau^K18^) or 12 hrs (Ub-αS) and diluted for immediate single-molecule measurement. Fluorescent protein samples were diluted to 100 pM (Ub-αS) or 40 pM (Ub-tau^K18^) final concentration and measured under flow according to previously established methods[30,38,39]. The bin time and flow rates for Ub-αS and Ub-tau^K18^ constructs were individually optimized to achieve the highest value of *Q* as described previously[12,30,38]. Ub-αS aggregates were measured at 100 µl/hr flow rate and the fluorescence signals were collected with 0.1 ms bin time. For Ub-tau^K18^ samples the flow rate and bin time were 50 µl/hr and 0.2 ms, respectively. Data were typically collected for 15 min at 25°C in frames of 50,000 bins. Independent triplicate experiments were performed for Ub-tau^K18^ and Ub-αS aggregation reactions. A representative set of aggregate degradation measurements is shown in Figure 3d and **e**.

### Single-molecule data analysis

We used the AND criterion to detect coincidence events in the two channels[31]. This separates aggregation events from background monomers by accepting only those signals for which the blue- and the red-excited channels are above the threshold value. The proportion of monomers that are associated to form oligomers is expressed using the association quotient *Q*, which is defined as

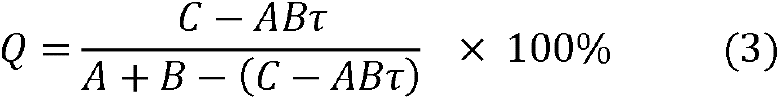

where *A* and *B* are the rates of detection of events in the two fluorescent channels, respectively, and *C* is the detection rate of coincident events; τ is the interval time of detection so that the *AB*τ expresses the coincident events that occur by chance[31]. We also measured the level of background detection of the buffer containing no fluorescent proteins and applied a uniform threshold five-fold over the background in both red- and blue-detector channels (10 kHz) to remove noise signals. A reference duplex DNA sample of 40 base pairs labeled with AlexaFluor 488 and AlexaFluor 647 at the 5’ end of each of the single DNA strands, repeatedly gave a *Q* value of 30%, using the same analysis[37].

Each single-molecule measurement was normalized to a standard number of frames and the number of significant coincident events was counted. The percentage decrease in oligomers upon proteasome treatment is calculated as follows:

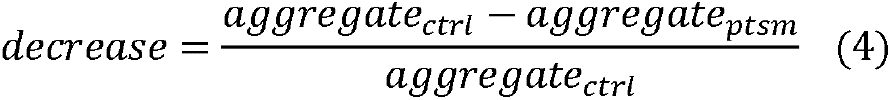

expressed in percentage. The estimation of *apparent aggregate size* was carried out as previously described[29]. In brief, the approximate monomer number per aggregate can be extracted assuming that 50% of monomers are donor using the equation

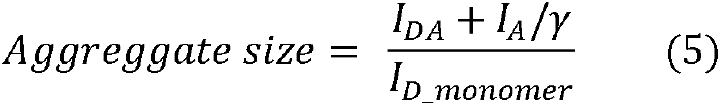

where *I_DA_* represents the donor fluorescence intensity in presence of acceptor, *I_A_* the acceptor fluorescence intensity. *I_D_monomer_* corresponds to the average intensity of donor monomers, and γ to a correction factor that accounts for different quantum yields and detection efficiencies of the donor and acceptor.

