## Supplementary material for "N-terminal ubiquitination of amyloidogenic proteins triggers removal of their oligomers by the proteasome holoenzyme": SFigures 1-5; STable 1

### Supplementary Figure 1

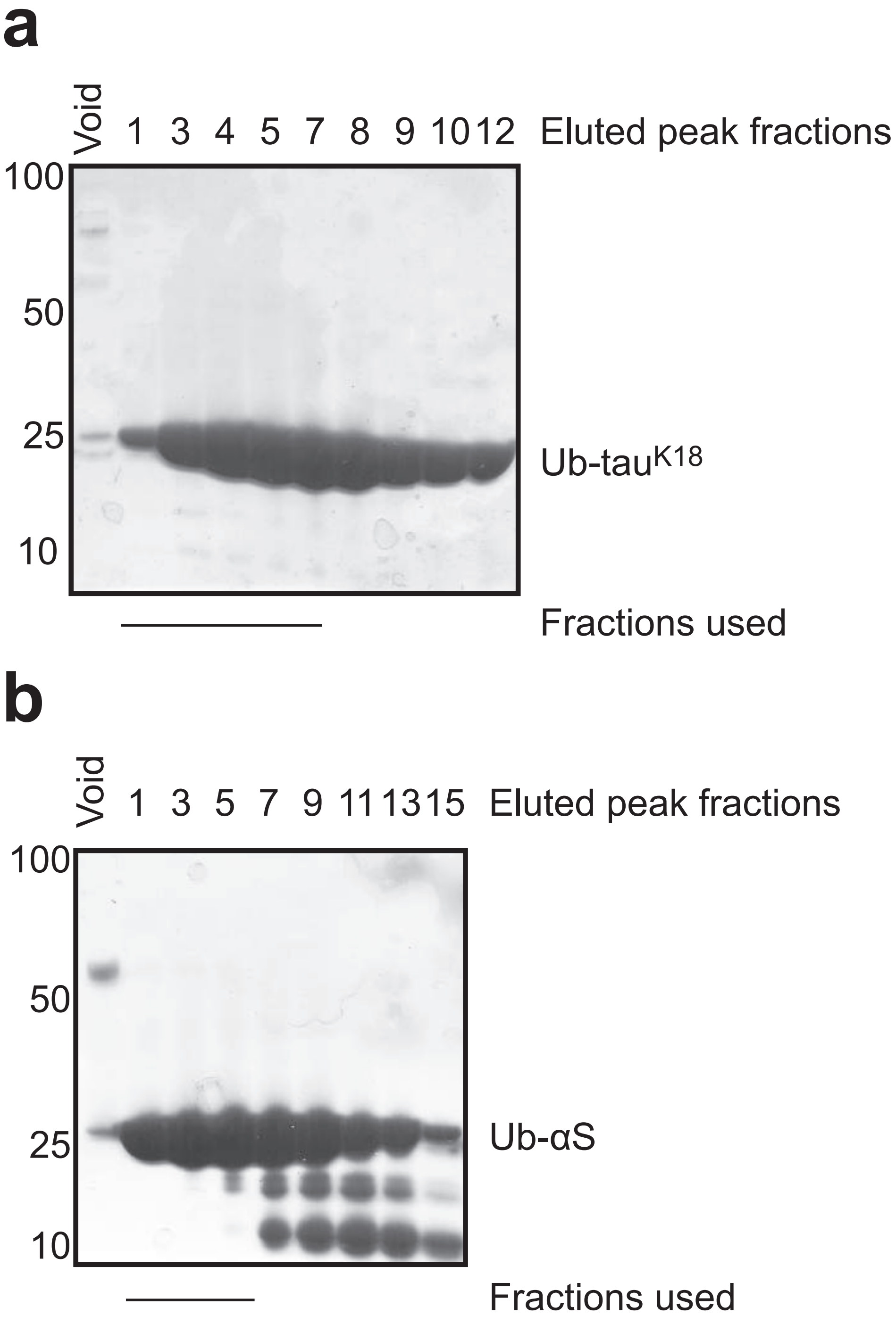

### Supplementary Figure 2

**a**

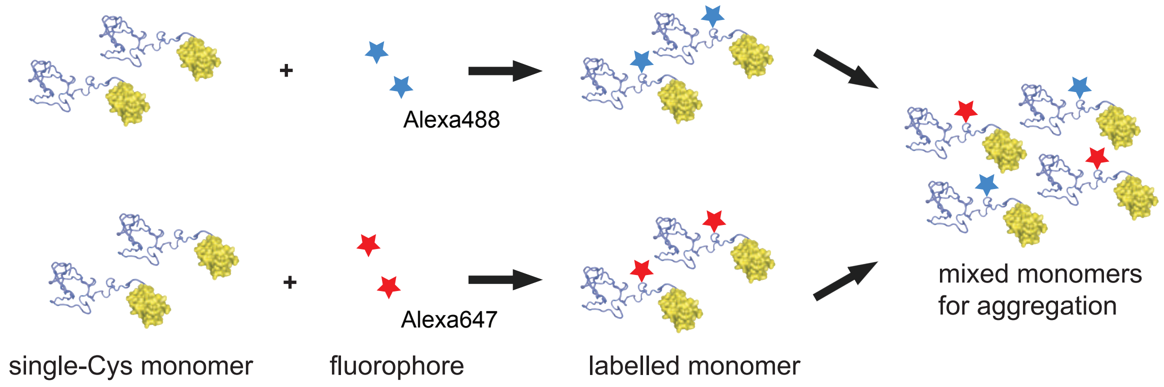


**b**

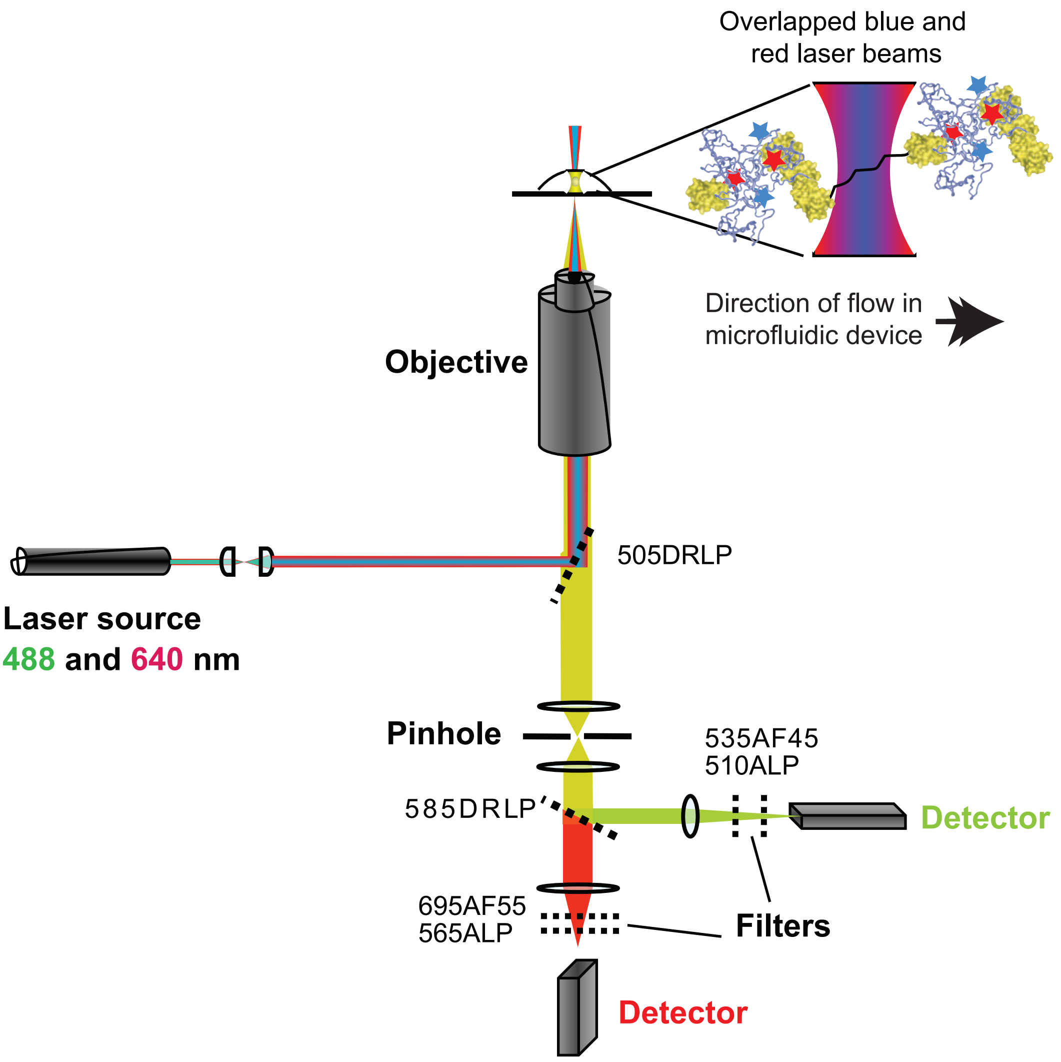

### Supplementary Figure 3

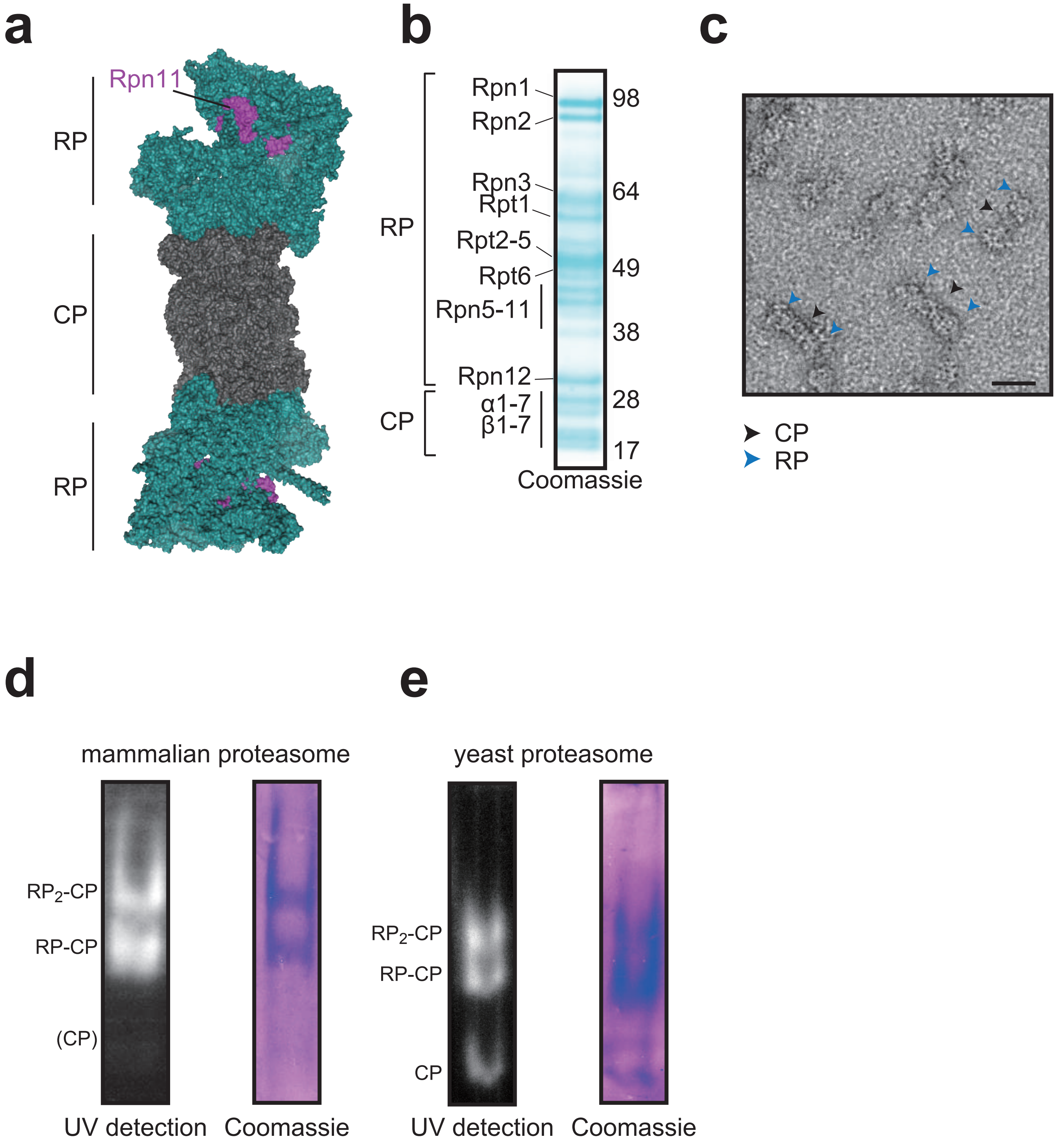

### Supplementary Figure 4

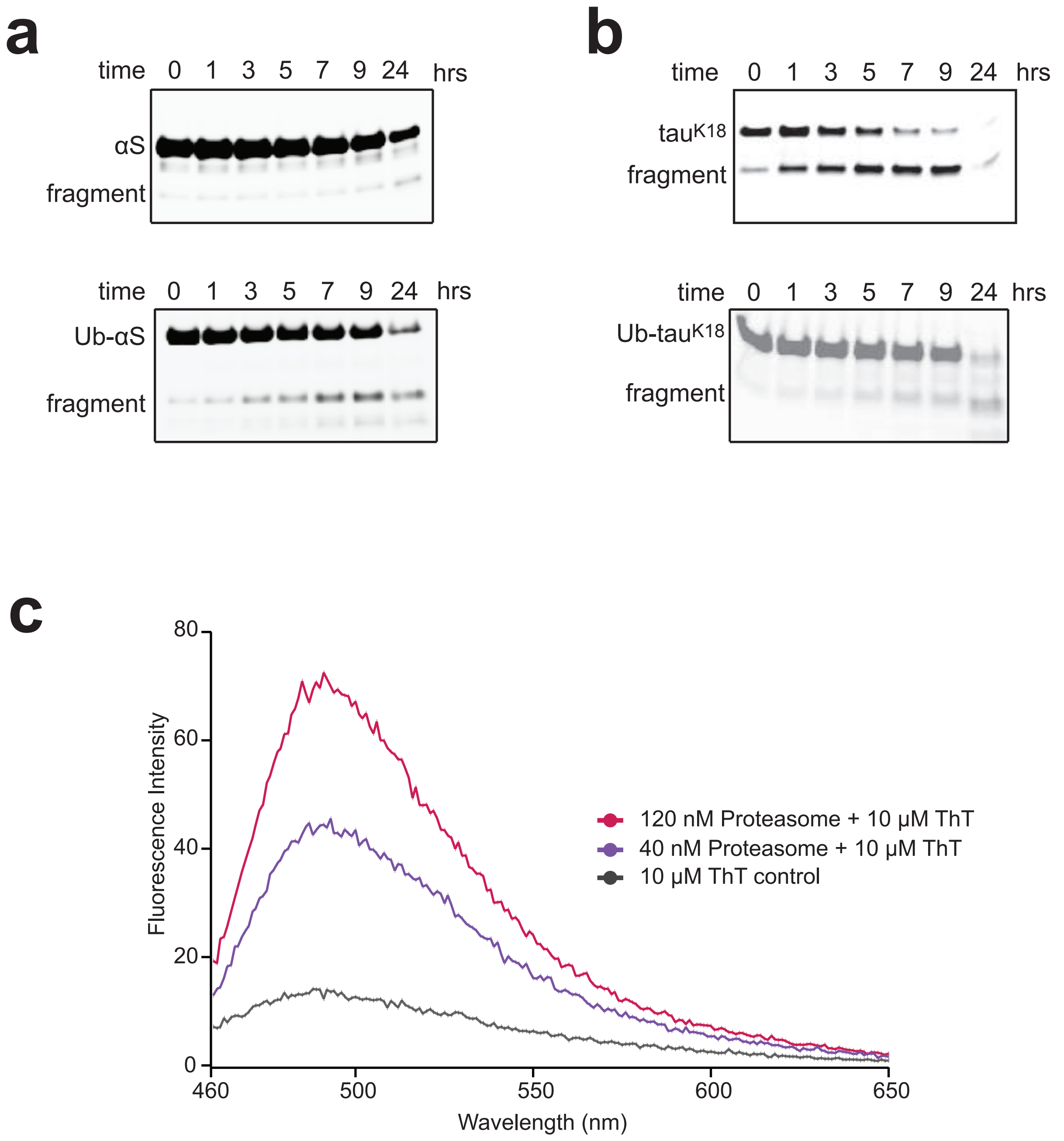

### Supplementary Figure 5

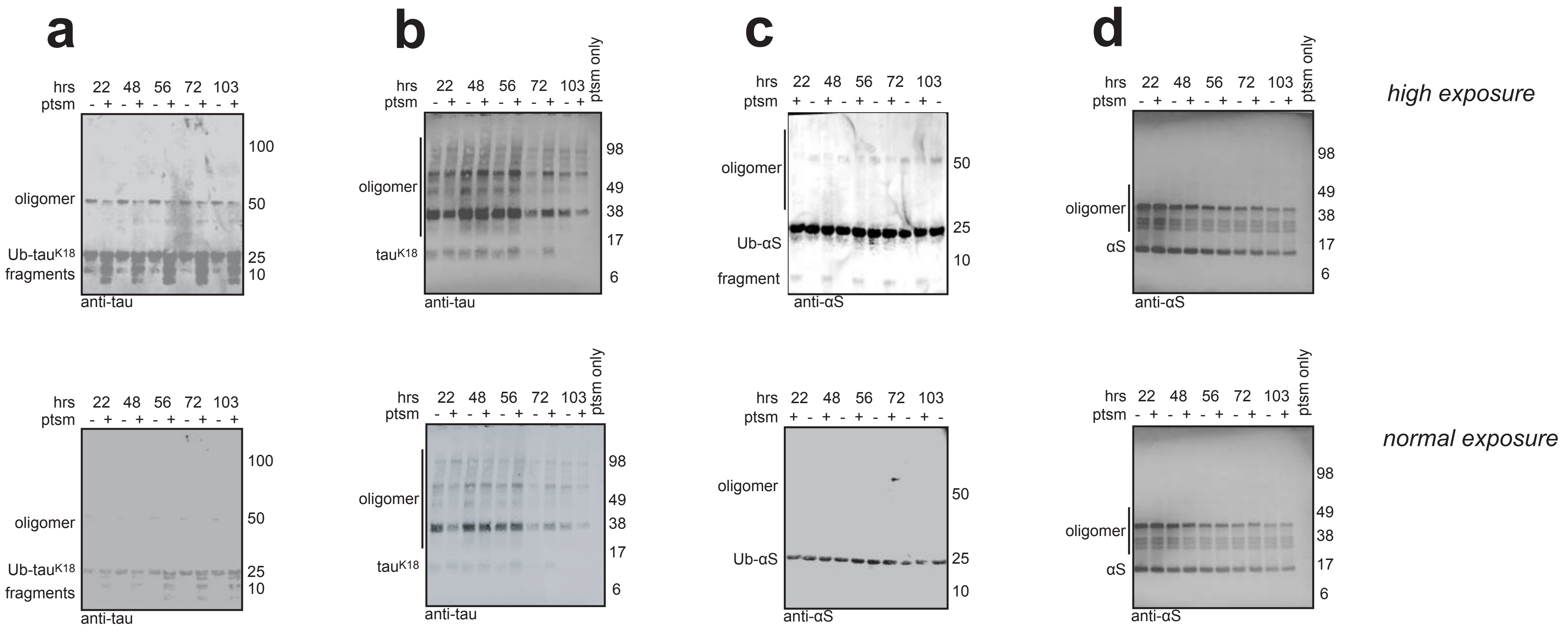

### Supplementary Table 1

#### a Size dependency of Ub-tau$^{K18}$ aggregates removed by the proteasome

| Aggregates | 4 hrs aggregation | | | 24 hrs aggregation | | | 48 hrs aggregation | | | 72 hrs aggregation | | | 96 hrs aggregation | | |
|---|---|---|---|---|---|---|---|---|---|---|---|---|---|---|---|
| | Control | +Proteasome | degraded | Control | +Proteasome | degraded | Control | +Proteasome | degraded | Control | +Proteasome | degraded | Control | +Proteasome | degraded |
| **Small** | 1023 | 57 | **94%** | 9284 | 519 | **94%** | 5234 | 292 | **94%** | 2624 | 1542 | **41%** | 272 | 94 | **65%** |
| **Medium** | 24 | 0 | **100%** | 456 | 36 | **92%** | 374 | 16 | **96%** | 224 | 116 | **48%** | 4 | 4 | **0%** |
| **Large** | 6 | 0 | **100%** | 62 | 6 | **90%** | 62 | 10 | **84%** | 32 | 18 | **44%** | 2 | 0 | **100%** |

#### b Size dependency of Ub-αS aggregates removed by the proteasome

| Aggregates | 4 hrs aggregation | | | 24 hrs aggregation | | | 48 hrs aggregation | | | 72 hrs aggregation | | | 96 hrs aggregation | | |
|---|---|---|---|---|---|---|---|---|---|---|---|---|---|---|---|
| | Control | +Proteasome | degraded | Control | +Proteasome | degraded | Control | +Proteasome | degraded | Control | +Proteasome | degraded | Control | +Proteasome | degraded |
| **Small** | 6230 | 1512 | **76%** | 2898 | 652 | **78%** | 2120 | 812 | **62%** | 1677 | 912 | **46%** | 3474 | 1765 | **49%** |
| **Medium** | 242 | 70 | **71%** | 30 | 8 | **73%** | 38 | 24 | **37%** | 54 | 34 | **37%** | 238 | 86 | **64%** |
| **Large** | 8 | 0 | **100%** | 2 | 0 | **100%** | 0 | 4 | **N/A** | 6 | 0 | **100%** | 14 | 24 | **N/A** |
